# Generating weighted and thresholded gene coexpression networks using signed distance correlation

**DOI:** 10.1101/2021.11.15.468627

**Authors:** Javier Pardo-Diaz, Philip S. Poole, Mariano Beguerisse-Díaz, Charlotte M. Deane, Gesine Reinert

## Abstract

Even within well-studied organisms, many genes lack useful functional annotations. One way to generate such functional information is to infer biological relationships between genes or proteins, using a network of gene coexpression data that includes functional annotations. Signed distance correlation has proved useful for the construction of unweighted gene coexpression networks. However, transforming correlation values into unweighted networks may lead to a loss of important biological information related to the intensity of the correlation. Here introduce a principled method to construct *weighted* gene coexpression networks using signed distance correlation. These networks contain weighted edges only between those pairs of genes whose correlation value is higher than a given threshold. We analyse data from different organisms and find that networks generated with our method based on signed distance correlation are more stable and capture more biological information compared to networks obtained from Pearson correlation. Moreover, we show that signed distance correlation networks capture more biological information than unweighted networks based on the same metric. While we use biological data sets to illustrate the method, the approach is general and can be used to construct networks in other domains.

**Data and code availability:** https://github.com/javier-pardodiaz/sdcorGCN

## 1 Introduction

Gene expression is the process by which genes in different organisms are *activated* to produce proteins when they are needed to carry out their function. Data relating to gene expression data contains key information about intracellular biological processes [11]. Gene coexpression datasets typically describe the expression level of different genes across different samples often taken under different experimental conditions. Such data are frequently represented as gene coexpression networks, with nodes representing genes and edges representing correlations in expression between pairs of genes across multiple samples [13]. Representing gene coexpression as networks helps in the study and visualisation of the expression data and the exploitation of the structure of interactions between genes at a whole-system level [27, 15]. One motivation behind creating these networks is that genes which are highly coexpressed across multiple samples are likely to have related functions [8, 21, 25, 16], allowing inference of gene function using *guilt by association* approaches [28]. This procedure is especially useful if the studied organism is poorly annotated. However, lack of reliable genomic information can hinder the validation of the accuracy of the network models generated from gene expression data. Noisiness in the data makes it difficult to distinguish genes that are expressed at low level from those not expressed. Therefore, there is a need for network construction pipelines that are robust to experimental error, yet capture most of the information in the data.

Multiple strategies to construct gene coexpression networks from gene expression data are available [1, 9, 24, 12, 7]. A recent novel approach uses signed distance correlation [17]. This approach identifies coexpression relationships between genes and produces robust networks that capture more biological information than those obtained using alternative metrics such as Pearson correlation, Spearman correlation and mutual information [17]. Signed distance correlation is based on distance correlation [22], a measure that evaluates an association between the pattern of changes in the samples, and provides a non-negative score that is zero if and only if the expression vectors are statistically independent. Because it is unsigned, distance correlation values do not permit the differentiation of genes with the same expression pattern and genes expressed at opposite times. Signed distance correlation overcomes this problem combining the distance correlation value with a sign that indicates the direction of the association. The sign corresponds to the sign of the Pearson correlation between the expression of the two genes across the samples in the dataset. Figure 1 below illustrates how signed distance correlation is obtained.

**Figure 1:**
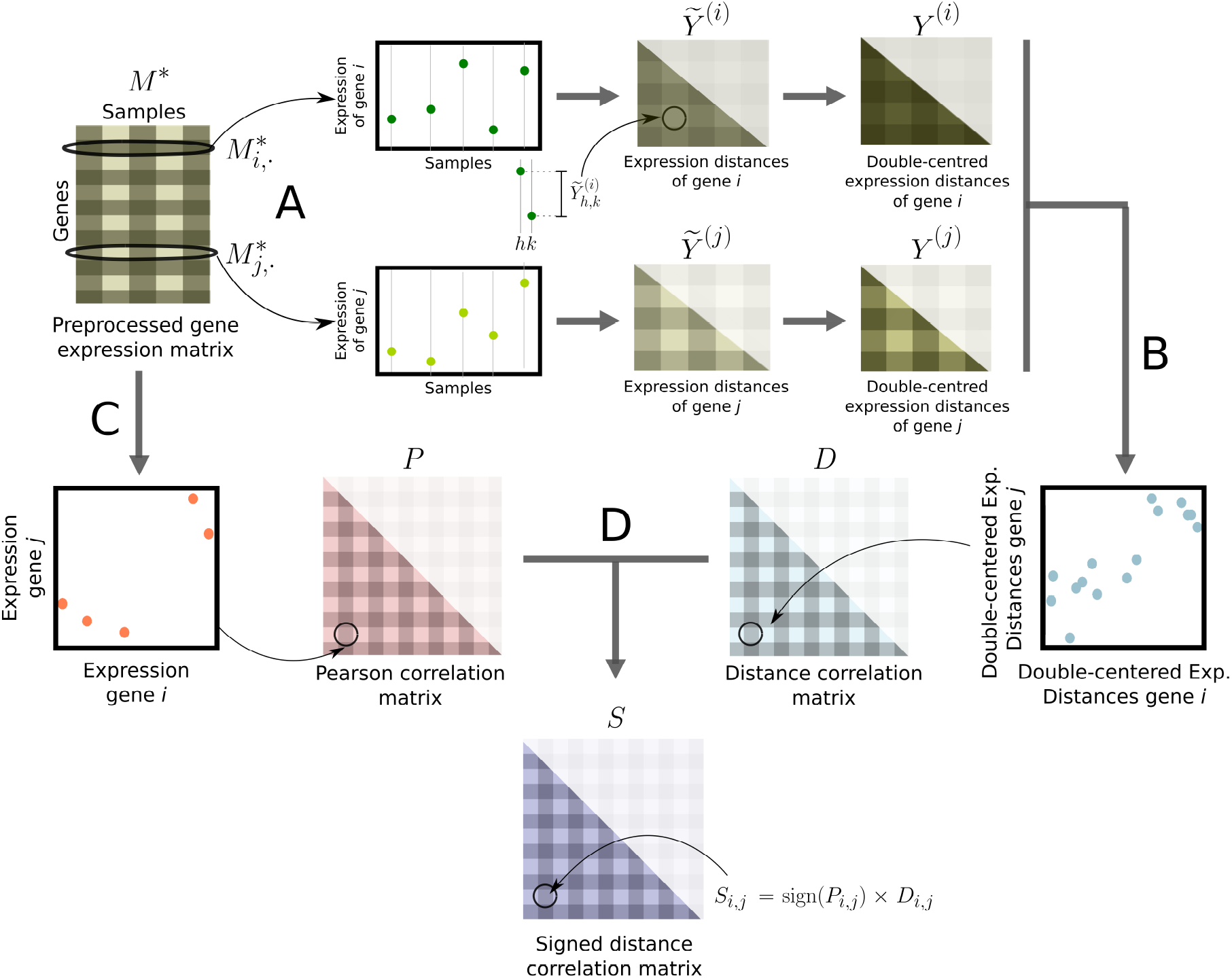
Pipeline to construct networks from gene expression data using signed distance correlation, adapted from [17]. **A**: We compute the expression distance matrices 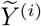 and 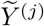 for each gene *i, j* ∈ {1, …, *m*}, and we double center them to obtain *Y*^(*i*)^ and *Y*^(*j*)^. **B**: We compute the distance correlation matrix *D*, whose entries *D_i,j_* are the positive root of the Pearson correlation between *Y*^(*i*)^ and *Y*^(*j*)^, for every pair of genes. **C**: We compute the Pearson correlation between each pair of rows in the *M*\* to obtain the Pearson correlation matrix *P*. **D**: To construct the signed distance correlation matrix *S* we multiply every distance correlation between the expression of two genes *D_i,j_* by the sign of their Pearson correlation sign(*P_i,j_*).

To our knowledge, signed distance correlation has been used in this field only to generate unweighted networks, using the R package COGENT [4]. COGENT aids the selection of a robust network construction method without the need for any external validation data. The main functions in this package iteratively split the gene expression data in two different sets, construct a network from each of them and then evaluate their similarity; the higher the similarity of the networks, the more robust the construction method. The functions implemented in COGENT to measure the similarity between the two generated networks are suitable to analyse only unweighted networks and complete weighted networks.

COGENT assists the selection of the optimal threshold value so that only pairs of genes for which the correlation value of their expression exceeds the threshold are connected in the network. The use of a threshold to construct networks results in a relatively sparse unweighted network. However, unweighted networks obtained through thresholding weighted edges may ignore important information about the strength of the correlation. Avoiding the use of a threshold results in complete weighted networks in which the weights of the edges are proportional to the correlation values. In this type of networks, it may be difficult to distinguish the signal from the noise. Thus, there is a need for a method that eases the signal and noise separation while keeping information about the strength of the correlation.

In this paper, we present a method to construct weighted and thresholded gene coexpression networks in which the sparsity can be controlled by assigning weighted edges only to those pairs of genes with an expression correlation higher than a given threshold. The weights of the edges correspond to the correlation values. To select the threshold value we have extended COGENT, including a comparison methodology that allows us to evaluate the similarity between weighted and thresholded networks.

Using this extension we construct and compare networks constructed from gene expression data using Pearson or signed distance correlation. For ease of comparison and for breadth, we analyse the three datasets presented in [17] as they not only correspond to different organisms (bacteria, yeast and human), but also are derived from three different experimental techniques: microarrays, RNA-Seq and single-cell RNA-Seq. When comparing the networks, we evaluate the robustness measurement from COGENT and the amount of biological information they capture according to STRING, a proteinprotein interaction database with scores for pairs of proteins [23]. The higher the STRING score for a protein pair, the more likely the pair is to have a biologically meaningful functional relationship. Using STRING, we show that for our datasets, networks constructed using signed distance correlation capture even more biological information and are structurally more stable than networks based on Pearson correlation.

We also compare the resulting weighted and thresholded networks based on signed distance correlation to the unweighted networks obtained using the same metric as presented in [17]. In this comparison, the weighted and thresholded networks capture more biological information than the corresponding unweighted networks, according to STRING.

While we apply our method to gene expression data, our method to construct networks from signed distance correlations (in combination with COGENT) can be used in applications beyond gene expression and beyond bioinformatics.

Data and source code are available from https://github.com/javier-pardodiaz/sdcorGCN and http://opig.stats.ox.ac.uk/resources.

## 2 Methodology

### 2.1 Datasets and pre-processing

We analyse the three datasets employed in [17]:

- RL3841: A collection of 54 microarrays measuring the expression of 7,077 genes of the bacterium *Rhizobium leguminosarum* bv. *viciae* 3841.
- Yeast: A dataset obtained using RNA-Seq of yeast (*Saccharomyces cerevisiae*) expressing pathways designed to increase ATP or GTP consumption. We use all the raw-counts for experiment E-MTAB-5174 in Expression Atlas [18] and remove the genes with zero expression variance. The final dataset which we feed into our pipeline includes the expression of 6,930 genes across 209 samples.
- Human liver: A dataset obtained using single-cell RNA-Seq of human liver cells [10]. The original dataset measures the expression of 15,353 genes in 1,622 cells.

The three datasets in the form of expression matrices (genes in rows and samples in columns) are publicly available online. We denote the expression matrices by *M*.

As detailed in [17], for the RL3841 and yeast datasets, we apply quantile normalisation [3] to the gene expression matrix *M*. This normalisation enables us to compare data from different experiments. To avoid interference from low expression values in the quantile normalisation, we ignore the 20% least expressed genes from each sample before the normalisation step. After the quantile normalisation, we set the ignored values to the lowest expression value in *M* to decrease the level of noise.

For the human liver dataset, as described in [17], we follow a different approach due to differences in the data – the gene expression levels in the dataset correspond to the expression of the genes different cells instead of to different samples – and the organism – while both *R. leguminosarum* and *S. cerevisiae* are unicellular organisms, humans are not. Hence, a considerable proportion of genes are not expressed in the studied cells. For this dataset, we quantile-normalise the data [3] to make the measurements in the different cells comparable. Afterwards, as in [19], we identify the *non-changing genes*. These genes are those for which the difference between its highest and lowest expression value (“expression difference”) is lower than the median of all the expression differences calculated for each gene, and in addition for which the mean expression signal between samples is lower than the median of all the expression signals calculated for each gene. After removing the “non-changing genes” we obtain an already quantile-normalised dataset with information for 8,585 genes.

In all three datasets, we denote the pre-processed gene expression matrix by *M*\*. This matrix has *n* rows (genes) and *m* columns (samples).

### 2.2 Correlation matrices

From each pre-processed gene expression matrix *M*\* we compute two correlation matrices *S* and *P*, both of them with dimensions *n* × *n*. The matrix *S* contains the signed distance correlation values for the expression of each pair of genes, whereas the matrix *P* includes the pairwise Pearson correlation values. The matrix *S* is the result of assigning the sign of the values in the matrix *P* to the distance correlation values [22] between the expression of the genes. Assigning a sign to the (unsigned) distance correlation values allows us to differentiate positive and negative correlation values. The pipeline followed to construct the expression matrices is described in [17] and depicted in Figure 1.

### 2.3 Weighted and thresholded network construction

In each of our three datasets, we use the gene correlation matrices *S* and *P* to obtain weighted and thresholded networks. These networks contain edges only between those pairs of genes that show a correlation of their expression greater than a given threshold. Unlike in unweighted networks, we assign a different weight to each edge. The weight of the edge corresponds to the value of the correlation of the expression between the two genes. The weighted and thresholded networks obtained using signed distance correlation matrix with threshold *ϕ* have an adjacency matrix *B_S_*(*ϕ*):

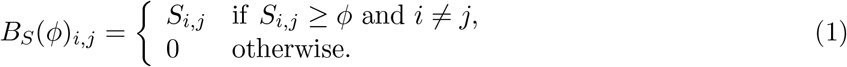

We construct the weighted and thresholded networks from the Pearson correlation matrix *P* in a similar way.

We use an extension of COGENT [4] to find the threshold value *ϕ*\* that results in a network with optimal self-consistency. The main idea behind COGENT is that the more similar two networks obtained using the same network construction method and from different overlapping subsets of the same dataset, the more self-consistent the employed method is: despite the changes in the dataset, the topology of the resulting networks is similar. Heuristically, the networks obtained using a self-consistent construction method are self-consistent themselves since small changes in the datasets will not affect their structure. Our contribution to COGENT relies on the addition of a method that allows assessing the similarity between weighted and thresholded networks (Equation 2).

As in [17], we run 25 COGENT iterations in which the samples (columns) in *M* are grouped into two overlapping sets *M*_1_ and *M*_2_. Both sets share half of the total number of samples, and differ in 1/3 from each other. At each iteration, we generate 25 pairs of correlation matrices *S*_1_ and *S*_2_ and test different threshold values. For each iteration and threshold value, the correlation matrices *S*_1_ and *S*_2_ are thresholded (Equation 1) and turned into networks *H*_1_ and *H*_2_. The processes taking place in each of the COGENT iterations to obtain the networks *H*_1_ and *H*_2_ are depicted in Figure 2. To calculate a similarity between the two networks *H*_1_ and *H*_2_, we use an adjusted weighted Jaccard index that permits the comparison between networks with different edge densities:

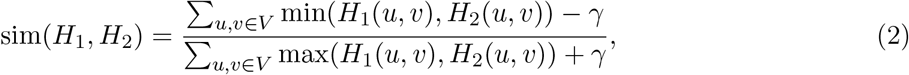

where

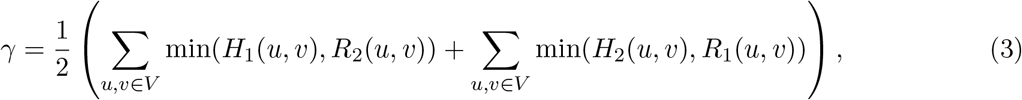

*R*_1_ and *R*_2_ are randomisations of the adjacency matrices *H*_1_ and *H*_2_, respectively, and *V* is the set of nodes in the network. To generate the randomisations, we first permute the rows in the correlation matrices and then shuffle the columns using the same permutation. The networks *R*_1_ and *R*_2_ have the same topological properties than *H*_1_ and *H*_2_ and allow us to estimate the expected similarity between the analysed networks and random networks with the same edge weight distribution. Without this correction, the networks derived from low threshold values would have the highest similarity values.

**Figure 2:**
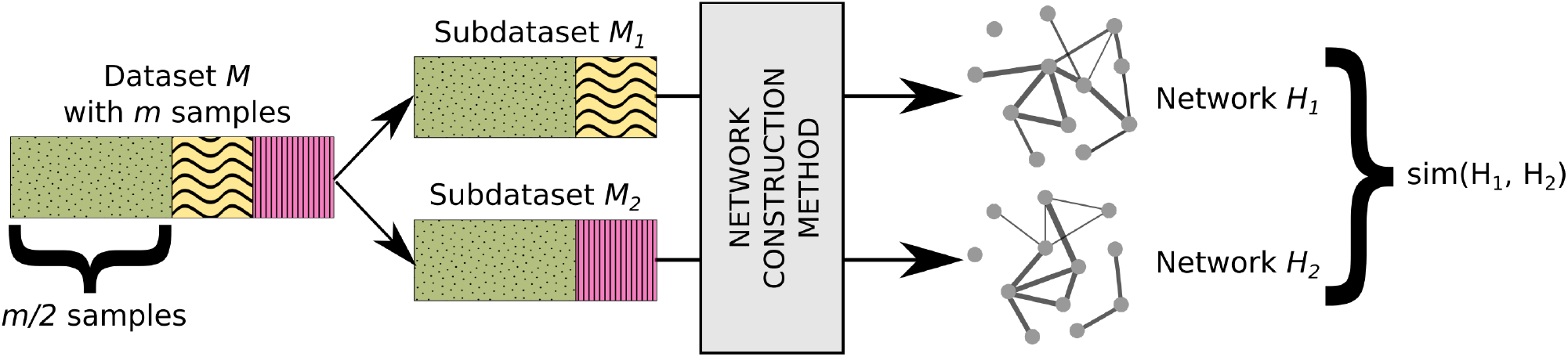
Illustration of a COGENT iteration. At each iteration, from the full dataset *M* with all *m* samples we generate two subdatasets *M*_1_ and *M*_2_. Each of the subdatasets contains half of the samples included in *M* (repesented in green with dots in the image). The rest of the samples in *M* are equally divided between *M*_1_ (yellow with waves) and *M*_2_ (pink with stripes), resulting in the two datasets having the same number of samples. Afterwards, from each dataset we construct gene coexpression network (network *H*_1_ from subdataset *M*_1_ and network *H*_2_ from subdataset *M*_2_) using the same network construction method, which might include pre-processing steps. Lastly, we obtain a similarity measure based on the overlap between the networks *H*_1_ and *H*_2_.

The similarity score associated with each threshold *s*(*ϕ*) is the average of the similarity of 25 pairs of networks (Equation 2). Then, to favour signal over noise we prioritise those networks with a low sum of edge weights and obtain the Score(*ϕ*) value associated to each threshold *ϕ*:

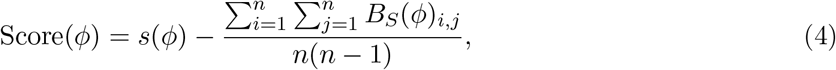

where *n* denotes the number of nodes in the network (i.e. the number of genes). After computing the scores for all tested thresholds, we select the threshold value *ϕ*\* that results in the highest score Score(*ϕ*\*). We denote the optimal weighted and thresholded network obtained using signed distance correlation as *WS*(*w_S_*). We denote the optimal sum of edge weights for the signed correlation by *w_S_*.

Following the same pipeline, we select the unique threshold value *ϕ*^⋆^ that results in the optimal network *WP*(*w_P_*) (sum of edge weights = *w_P_*) obtained using the Pearson correlation matrix *P*.

### 2.4 STRING evaluation of weighted and thresholded networks

We assess the amount of biological information contained in the just generated weighted and thresh-olded networks *WS*(*ws*) and *WP*(*w_P_*) using STRING, a database of known and predicted protein–protein interactions [23]. STRING collects information from numerous sources, including experimental data, computational predictions and textmining. The association evidence in STRING is categorized into independent channels, weighted, and integrated to produce a confidence score *C* for all recorded protein interactions. Interactions with high *C* score are more likely to be true than those with a low score. To evaluate the networks, we employ the same set of confidence scores as in [17]:

- *C*: Total scores provided by STRING.
- *C*^†^: Scores that *only* consider coexpression information
- *C*^‡^: Scores that *exclude* coexpression information.

We expect the overlap of our networks with the scores in *C*^†^ to be higher than the overlap with the scores in *C*^‡^ since our input datasets contain gene expression information. Nevertheless, both *C*^†^ and *C*^‡^ are interesting to analyse: the first one might be an indicator on how well the networks capture coexpression relationships whereas the latter can be used as in indicator on how well coexpression can predict other types of relationships.

For each network and confidence score set we compute the dot product of the edge weights and the STRING confidence score. Afterwards, we divide the result by the sum of the weights of all edges in the network. Equation 5 indicates how we obtain the STRING score for a weighted network with adjacency matrix *B_S_*(*ϕ*) and confidence score set *C*:

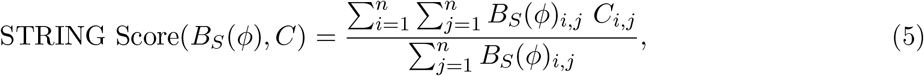

where *n* denotes the number of nodes in the network, *B_S_*(*ϕ*)_*i,j*_ is the the entry for genes *i* and *j* in *B_S_*(*ϕ*), and *C_i,j_* refers to the confidence score for the genes *i* and *j* in the confidence score set *C*.

Despite being normalised for the sum of edge weights, the score might be biased towards networks with fewer and heavier edges. For this reason, and in order to keep the comparison between the networks obtained using Pearson and distance correlation the fairest possible, we construct and evaluate six additional networks:

- *WS*(*w_P_*): Network from *S* with the same sum of weights of edges as *WP*(*w_P_*)
- *WS*(*e_P_*): Network from *S* with the same number of edges as *WP*(*w_P_*)
- *WS*(*a_P_*): Network from *S* with the same average edge weight as *WP*(*w_P_*)
- *WP*(*w_S_*): Network from *P* with the same sum of weights of edges as *WS*(*w_S_*)
- *WP*(*e_S_*): Network from *P* with the same number of edges as *WS*(*w_S_*)
- *WP*(*a_S_*): Network from *P* with the same average edge weights as *WS*(*w_S_*)

We also generate two sets of 30 random networks to compare the results obtained in the STRING evaluation with those expected by chance. The sets of random networks have the same number of edges and distribution of edge weights as *WS*(*w_S_*) and *WP*(*w_P_*), respectively. We generate the networks by assigning the edges of the original networks to randomly picked pairs of vertices as a Bernoulli graph with fixed number of edges ad weights, and compute their STRING scores.

### 2.5 Comparison of weighted and thresholded networks and unweighted networks

Next, we assess whether the weighted and thresholded gene coexpression networks we obtain capture more biological information than unweighted coexpression networks. We compare our networks with the optimal unweighted signed distance correlation networks from [17], denoted by *NS*(*d_S_*). We analyse and compare four networks:

- *WS*(*w_S_*) Optimal weighted network from *S*
- *NS*(*w_S_*) Unweighted network with the same edges as *WS*(*w_S_*)
- *NS*(*d_S_*) Optimal unweighted network from *S*
- *WS*(*d_S_*) Weighted network with the same edges as *NS*(*d_S_*)

We focus only on signed distance correlation networks because unweighted and weighted signed distance correlation networks are both more self-consistent and capture more biological information than their matching Pearson networks (see Results section).

Unweighted networks, by definition, do not have weights associated with their edges; in order to be able to compare the networks using the scoring function from Equation 5, we assign to edges of the networks *NS*(*w_S_*) and *NS*(*d_S_*) as weight the average edge weight of the networks *WS*(*w_S_*) and *WS*(*d_S_*), respectively. All the edges in an unweighted networks thus have the same weight.

## 3 Results

### 3.1 Network construction and COGENT evaluation

We evaluate how the score retrieved using COGENT changes across different thresholds in each dataset and for each correlation matrix. This score depends on the similarity of the networks constructed at each COGENT iteration. The similarity is adjusted using the overlap expected between each of the networks and a networks with the same edge weights distributed randomly, and then adjusted to prioritise sparser networks (see Equations 2 and 4). Figure 3 illustrates how the signed distance and Pearson scores change for different edge weight values in the three datasets. In all the datasets (RL3841, yeast, and human liver) the curve shows a similar shape and there is an edge weight value for which the scores reach their maxima. The thresholds associated with those edge weights are the optimal thresholds *ϕ*\* and *ϕ*^⋆^ which we choose when constructing the networks *WS*(*w_S_*) and *WP*(*w_P_*). The score, threshold and summaries of these networks are shown in Table 1. In all cases, the highest scores obtained using signed distance correlation are higher than those for Pearson correlation. Therefore, weighted and thresholded gene coexpression networks based on signed distance correlation can be more self-consistent than those based on Pearson correlation.

**Figure 3:**
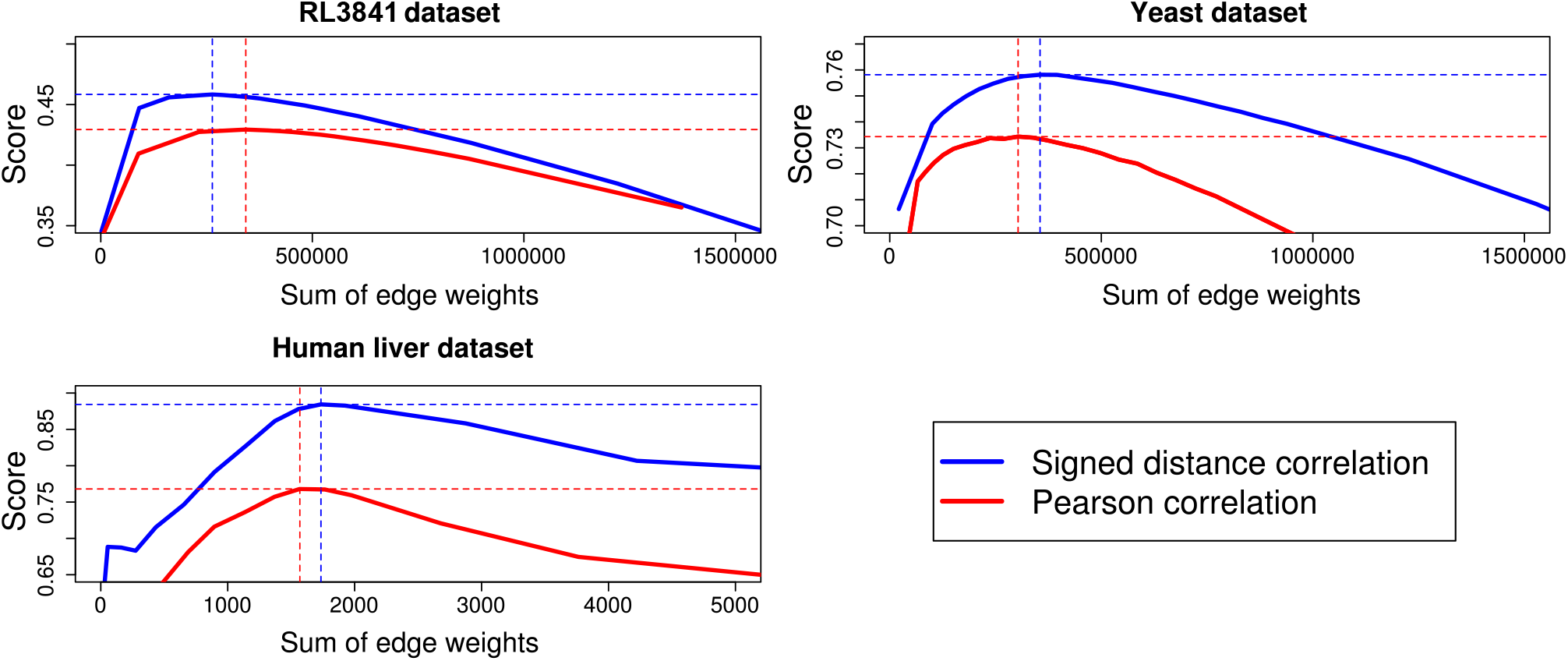
Self-consistency scores for different thresholds and datasets. The blue and red lines show the scores (Equation 4) of networks obtained using signed distance and Pearson correlations, respectively, to generate weighted and thresholded networks. The dashed vertical and horizontal lines indicate the optimal sums of edge weights and the highest scores, respectively. Scores obtained by signed distance correlation are almost uniformly higher than scores from Pearson correlation.

**Table 1:** Metrics and COGENT score for the optimal networks of the studied datasets. “Corr” indicates the correlation matrix employed and “Thr” denotes optimal threshold

| RL3841 dataset |  |  |  |  |  |  |
| --- | --- | --- | --- | --- | --- | --- |
| Network | Corr | COGENT score | Thr | Number of Edges | Sum of weights | Average weight |
| $WS(w_S)$ | $S$ | 0.46 | 0.60 | 388374 | 263662 | 0.6789 |
| $WP(w_P)$ | $P$ | 0.43 | 0.55 | 531374 | 342785 | 0.6451 |
| Yeast dataset |  |  |  |  |  |  |
| Network | Corr | COGENT score | Thr | Number of Edges | Sum of weights | Average weight |
| $WS(w_S)$ | $S$ | 0.76 | 0.82 | 405576 | 355227 | 0.8759 |
| $WP(w_P)$ | $P$ | 0.73 | 0.81 | 350549 | 303310 | 0.8652 |
| Human liver dataset |  |  |  |  |  |  |
| Network | Corr | COGENT score | Thr | Number of Edges | Sum of weights | Average weight |
| $WS(w_S)$ | $S$ | 0.88 | 0.44 | 3320 | 1734.58 | 0.5225 |
| $WP(w_P)$ | $P$ | 0.77 | 0.44 | 3047 | 1567.93 | 0.5146 |

### 3.2 STRING evaluation

We assess the amount of biological information contained in the networks following the methodology described in the Methods section. Tables 2–4 present the networks constructed using the different datasets, their metrics, and their STRING scores. The optimal signed distance correlation networks *WS*(*w_S_*) retrieve a higher score than all alternatives based on Pearson correlation for all the studied datasets. The optimal Pearson correlation networks *WP*(*w_P_*) retrieve a lower score than most of their (signed distance) competitors. The only exception for this trend is the yeast network *WS*(*a_P_*) (signed distance correlation network with the average edge weight of the optimal Pearson correlation network) which returns a lower score than the network *WP*(*w_p_*) (optimal Pearson correlation network). These results suggest that using signed distance correlation for generating weighted and thresholded networks from gene expression data can yield better results than with Pearson correlation. Figure 4 shows the STRING scores obtained by the networks with sum of edge weights *w_S_* and *w_P_* for the three datasets and the different sets of evidence.

**Table 2:** Metrics and STRING scores for the networks from the RL3841 dataset. RE indicates the expected result and its standard deviation. *C*, *C*^†^ and *C*^‡^ indicate the scores obtained for each of the different sets of confidence from STRING. For each set of networks, the highest score values are indicated in bold.

| Network | Corr | Number of Edges | Sum of weights | Average weight | $C$ | $C^\dagger$ | $C^\ddagger$ |
| --- | --- | --- | --- | --- | --- | --- | --- |
| $WS(w_S)$ | $S$ | 388374 | 263662 | 0.6789 | **28.66** | **8.70** | **25.72** |
| $WP(e_S)$ | $P$ | 388374 | 261723 | 0.6739 | 24.49 | 7.31 | 22.08 |
| $WP(w_S)$ | $P$ | 391691 | 263662 | 0.6731 | 24.43 | 7.28 | 22.03 |
| $WP(a_S)$ | $P$ | 367002 | 249153 | 0.6789 | 24.86 | 7.52 | 22.39 |
| RE $w_S$ | $S$ | 388374 | 263662 | 0.6789 | 7.51<br>$\pm 0.08$ | 0.90<br>$\pm 0.02$ | 7.11<br>$\pm 0.08$ |
| $WP(w_P)$ | $P$ | 531374 | 342785 | 0.6451 | 22.39 | 6.26 | 20.26 |
| $WS(e_P)$ | $S$ | 531374 | 347174 | 0.6534 | 25.64 | 7.27 | 23.12 |
| $WS(w_P)$ | $S$ | 523675 | 342785 | 0.6546 | **25.78** | **7.33** | **23.23** |
| $WS(a_P)$ | $S$ | 585747 | 377861 | 0.6451 | 24.78 | 6.88 | 22.37 |
| RE $w_P$ | $P$ | 531374 | 342785 | 0.6451 | 7.53<br>$\pm 0.06$ | 0.91<br>$\pm 0.02$ | 7.12<br>$\pm 0.05$ |

**Table 3:** Metrics and STRING scores for the networks from the Yeast dataset. RE indicates the expected result and its standard deviation. *C*, *C*^†^ and *C*^‡^ indicate the scores obtained for each of the different sets of confidence from STRING. For each set of networks, the highest score values are indicated in bold.

| Network | Corr | Number of Edges | Sum of weights | Average weight | $C$ | $C^\dagger$ | $C^\ddagger$ |
| --- | --- | --- | --- | --- | --- | --- | --- |
| $WS(w_S)$ | $S$ | 405576 | 355227 | 0.8759 | 106.40 | 75.61 | 62.25 |
| $WP(e_S)$ | $P$ | 405576 | 347541 | 0.8569 | 105.69 | 75.60 | 61.58 |
| $WP(w_S)$ | $P$ | 415223 | 355227 | 0.8556 | 104.48 | 74.52 | 61.06 |
| $WP(a_S)$ | $P$ | 286663 | 251077 | 0.8759 | **124.81** | **92.58** | **69.84** |
| RE $w_S$ | $S$ | 405576 | 355227 | 0.8759 | 13.94<br>$\pm 0.14$ | 4.77<br>$\pm 0.07$ | 11.83<br>$\pm 0.13$ |
| $WP(w_P)$ | $P$ | 350549 | 303310 | 0.8652 | 113.36 | 86.36 | 64.90 |
| $WS(e_P)$ | $S$ | 350549 | 309784 | 0.8837 | 114.12 | 82.30 | 65.67 |
| $WS(w_P)$ | $S$ | 342774 | 303310 | 0.8849 | **115.34** | **83.36** | **66.21** |
| $WS(a_P)$ | $S$ | 486198 | 420679 | 0.8652 | 97.45 | 67.98 | 58.16 |
| RE $w_P$ | $P$ | 350549 | 303310 | 0.8652 | 13.96<br>$\pm 0.13$ | 4.77<br>$\pm 0.06$ | 11.84<br>$\pm 0.13$ |

**Table 4:**
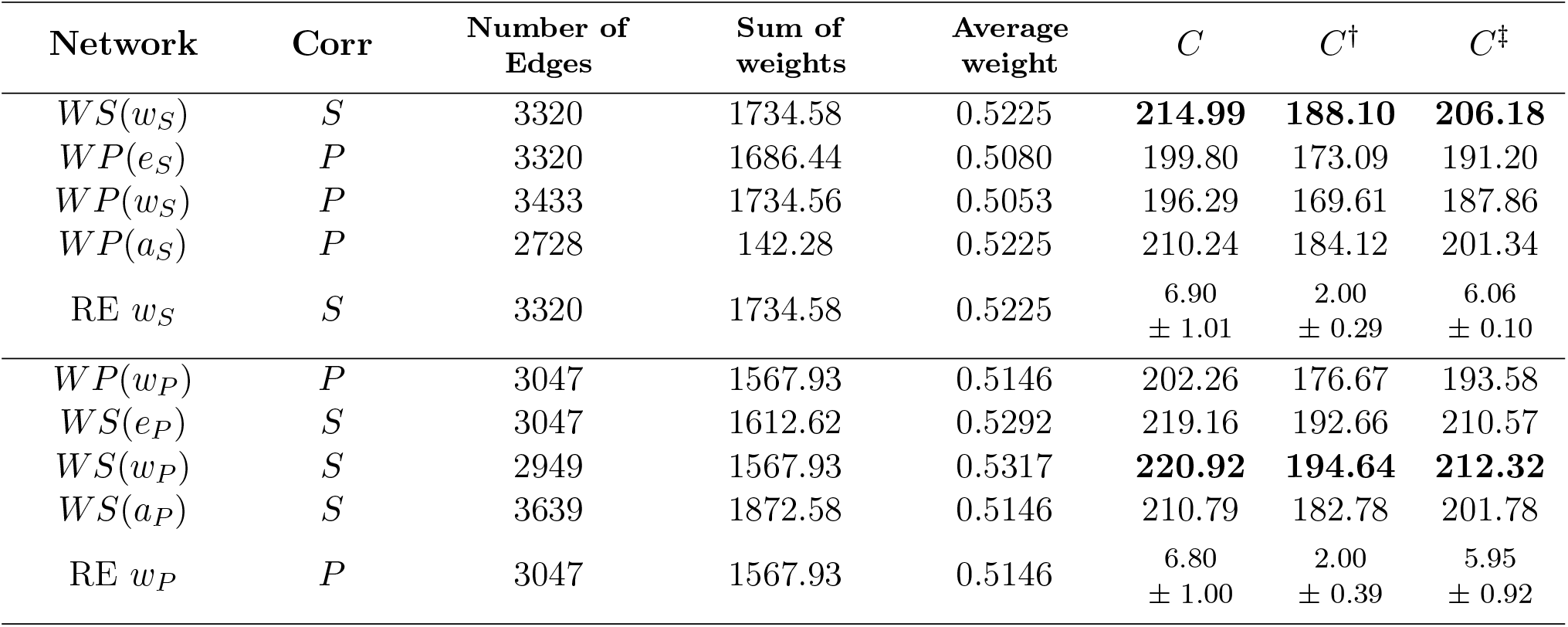
Metrics and STRING scores for the networks from the Human liver dataset. RE indicates the expected result and its standard deviation. *C*, *C*^†^ and *C*^‡^ indicate the scores obtained for each of the different sets of confidence from STRING. For each set of networks, the highest score values are indicated in bold.

**Figure 4:**
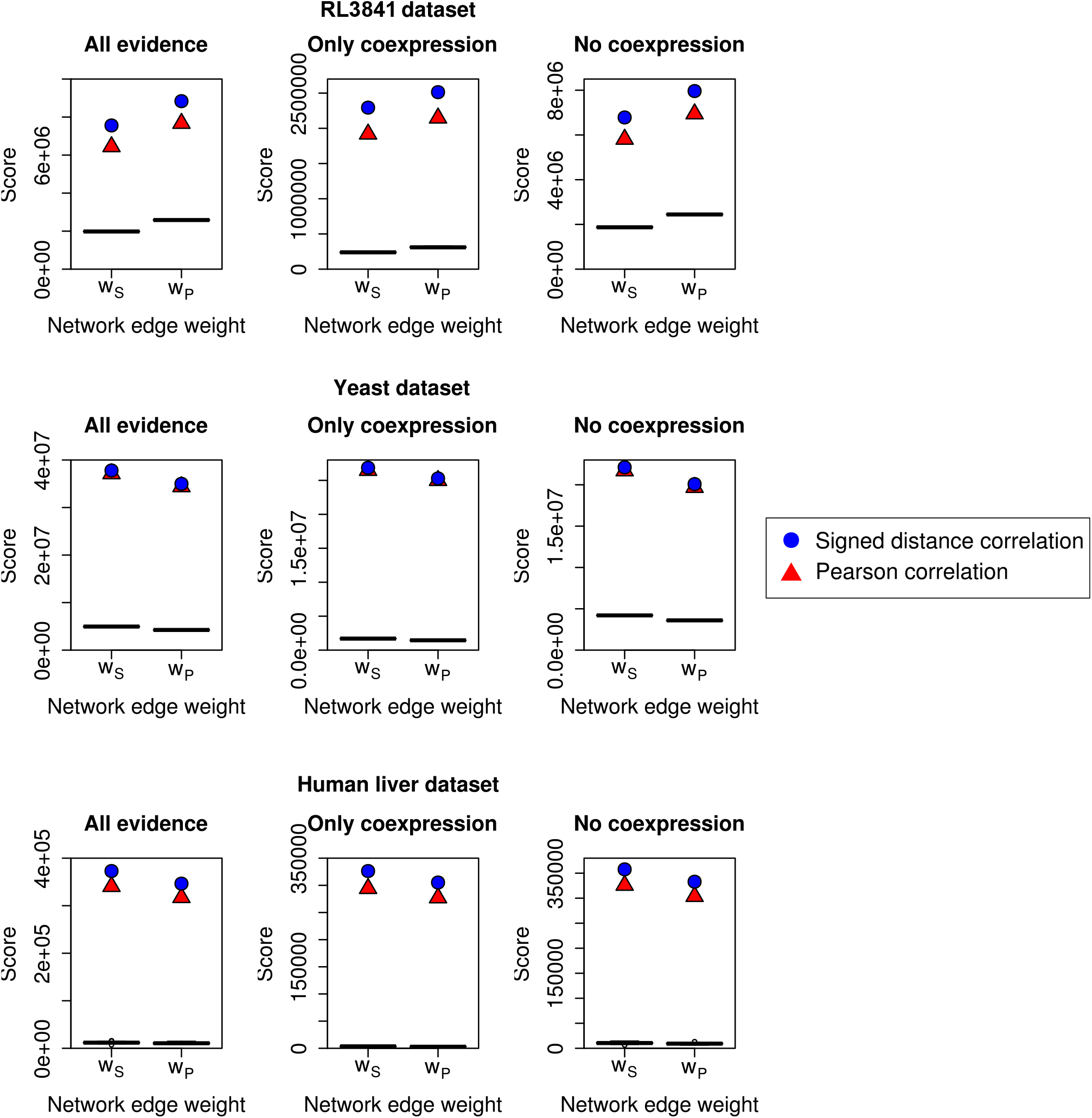
Scores obtained for the gene coexpression networks obtained from the three datasets using STRING. All panels show the score for the different networks in the y-axis, and the sum of weights the x-axis. The scores are the result of the dot product of the weights of the edges in the networks and the confidence scores with all evidence (*C*), only coexpression evidence (*C*^†^) and everything excluding coexpression (*C*^‡^) from STRING associated with them, divided by the sum of edge weights. The black box plots correspond to the scores obtained by 30 random networks. Blue circles and red triangles represent signed distance correlation and Pearson correlation, respectively.

We observe that for both correlation matrices, a higher average edge weight (and a lower number of edges) implies a higher STRING score. Figure 5 shows the high positive correlation between the STRING score using the information set *C* and the average edge weight of the networks for the RL3841 dataset. Still, often signed distance correlation networks retrieve a higher STRING score than Pearson networks even when their average edge weight is lower. For example, for the RL3841 dataset, the only case in which a Pearson correlation network retrieves a higher STRING score than a signed distance correlation network is when comparing the network *WS*(*a_P_*) (signed distance correlation network with the average edge weight of the optimal Pearson correlation network, with score 24.78) versus the network *WP*(*a_S_*) (Pearson correlation network with the average edge weight of the optimal signed distance correlation network, with score 24.86). However, this comparison lacks interest since their average edge weights (0.6789 and 0.6451) are quite different.

**Figure 5:**
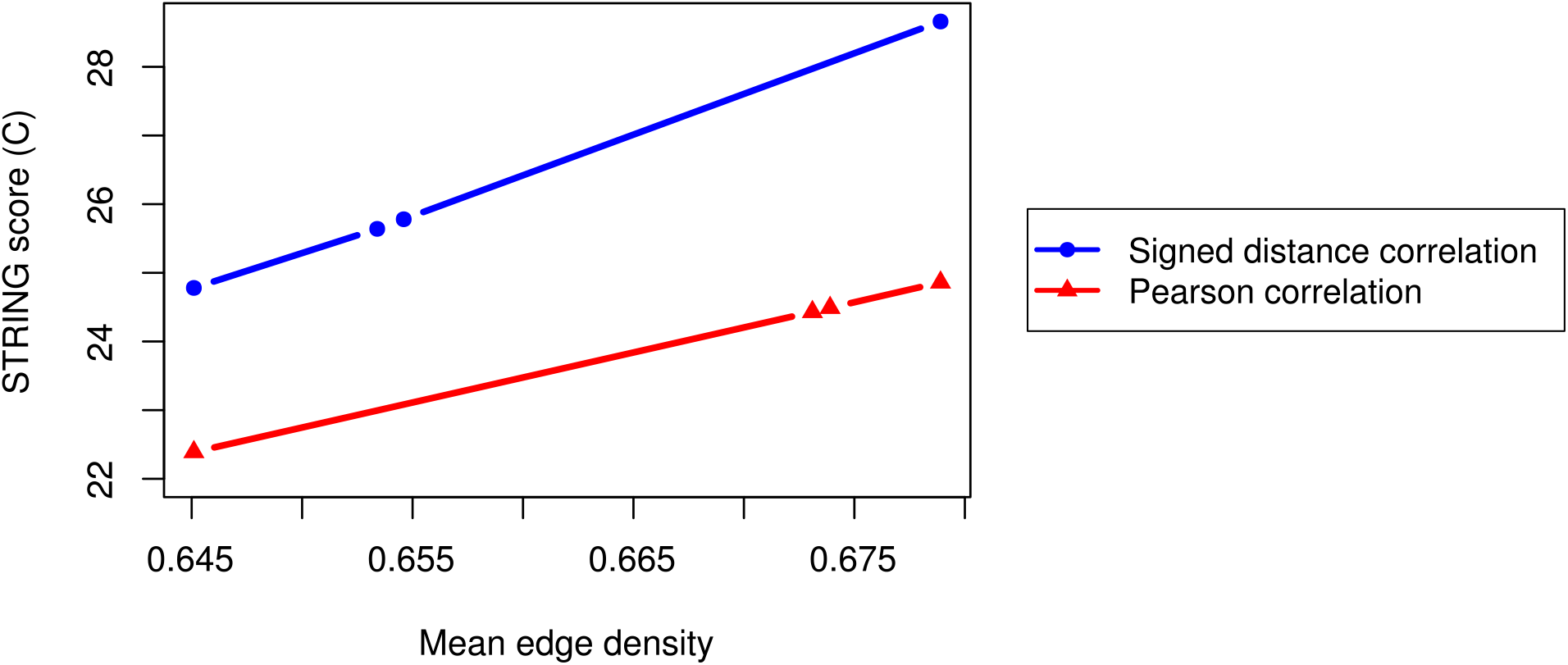
Distribution of STRING scores using the set of information *C* and average edge weight of the RL3841 networks. Blue and circles for signed distance correlation. Red and triangles for Pearson correlation

The random networks obtain a lower STRING score than Pearson and signed distance correlation networks (see Tables 2–4 and Figure 4). We find the highest difference between the random expected STRING scores and those retrieved by the networks when using only coexpression information (*C*^†^). The score obtained by the networks *WS*(*w_S_*) and *WP*(*w_P_*) are 9.63 and 6.89 times higher than the score expected by chance in the case of the RL3841 dataset. The smallest differences correspond to the exclusion of the coexpression information (*C*^‡^), where the values are 3.62 (*WS*(*w_S_*)) and 2.84 (*WP*(*w_P_*)) times higher than the randomly expected values for the RL3841 dataset.

### 3.3 Comparison with unweighted networks

Next we compare the amount of biological information captured by weighted and thresholded networks, and unweighted networks, constructed using signed distance correlation. We compare the two groups of networks with the same sets of edges as described in the Methods section: the optimal set of edges for the weighted and thresholded network and the optimal set for the unweighted network in [17]. We assign as weight to the edges in the unweighted networks the average weight in the matching weighted and thresholded networks. The results for the STRING evaluation for the different datasets are shown in Table 5. For the three datasets, the weighted and thresholded networks can capture more of the biological information and should therefore be preferred. For the human liver dataset we present only a set of two networks since the optimal threshold threshold for the weighted and the unweighted networks is the same and therefore they have the same edges.

**Table 5:** STRING scores for unweighted and weighted and thresholded networks for the three datasets using signed distance correlation. *C*, *C*^†^ and *C*^‡^ indicate the scores obtained for each of the different sets of confidence from STRING. Types “W” and “U” denote whether the networks are weighted or unweighted, respectively. Each pair of networks have the same edges; for each pair of networks, the highest score values are indicated in bold.

| RL3841 dataset |  |  |  |  |  |  |  |
| --- | --- | --- | --- | --- | --- | --- | --- |
| Network | Type | Number of Edges | Sum of weights | Average weight | $C$ | $C^\dagger$ | $C^\ddagger$ |
| $WS(w_S)$ | W | 388374 | 263662 | 0.6789 | **28.66** | **8.70** | **25.72** |
| $NS(w_S)$ | U | 388374 | 263662 | 0.6789 | 27.31 | 7.87 | 24.57 |
| $WS(d_S)$ | W | 313348 | 217914 | 0.6954 | **30.97** | **9.88** | **27.70** |
| $NS(d_S)$ | U | 313348 | 217914 | 0.6954 | 29.61 | 9.02 | 26.55 |
| Yeast dataset |  |  |  |  |  |  |  |
| Network | Type | Number of Edges | Sum of weights | Average weight | $C$ | $C^\dagger$ | $C^\ddagger$ |
| $WS(w_S)$ | W | 405576 | 355227 | 0.8759 | **106.40** | **75.61** | **62.25** |
| $NS(w_S)$ | U | 405576 | 355227 | 0.8759 | 103.49 | 72.96 | 61.12 |
| $WS(d_S)$ | W | 314771 | 279879 | 0.8892 | **120.12** | **87.59** | **68.19** |
| $NS(d_S)$ | U | 314771 | 279879 | 0.8892 | 117.36 | 85.03 | 67.14 |
| Human liver dataset |  |  |  |  |  |  |  |
| Network | Type | Number of Edges | Sum of weights | Average weight | $C$ | $C^\dagger$ | $C^\ddagger$ |
| $WS(w_S)$ | W | 3320 | 1734.58 | 0.5225 | **214.99** | **188.10** | **206.18** |
| $NS(d_S)$ | U | 3320 | 1734.58 | 0.5225 | 212.02 | 185.35 | 203.55 |

## 4 Discussion and conclusions

The method we propose combines the intuitiveness of the unweighted networks, as there are edges connecting only those pairs of genes with a high correlation in their expression, with the fine tuning provided by assigning different weight values to different edges. This combination allows to differentiate genes which are highly coexpressed from genes that even if they are coexpressed, their association is not high. These characteristics can help to study and analyse the networks.

We select the optimal threshold value to generate the weighted and thresholded networks using an extension of COGENT [4] that allows assessing the similarity of this type of networks. For each correlation matrix, we select the threshold value that results in a high self-consistency and prioritizes signal over noise. We also normalised the correlation values higher than the threshold between zero and one, using these normalised values as weights: edges connecting genes with a expression correlation less or equal to the threshold had a weight of zero; edges connecting genes with a expression correlation equals to one had a weight of one; edges connecting genes with a expression correlation between the threshold and one had weights between zero and one. We evaluated the self-consistency of the obtained network as described in Section 2.3. We observed that this normalisation step resulted in a similar self-consistency. For the signed distance correlation network, the optimal threshold was 0.52, resulting in a score of 0.4582. This value is lower than the one obtained without the normalisation: 0.4584. For the Pearson correlation networks, we also obtained higher values when not using the normalisation. In light of these results, we do not include this normalisation step in our pipeline.

The correlation matrices *S* and *P* differentiate positive from negative correlations. However, the threshold value we select to construct the networks is always greater than zero and therefore the weights of the edges in the network are always positive. As discussed in [17], we construct networks that include only positive correlations since negative correlations do not always imply a functional relationship. Nevertheless, a signed network with weights associated with their edges might include valuable information since the sign of the weights allow to differentiate positive and negative associations. Exploring a modification of our network construction pipeline of this type might provide an improvement.

For the three studied datasets, the use of signed distance correlation to generate weighted and thresholded networks results in a higher self-consistency than the use of Pearson correlation. Regarding the STRING evaluation, most of the times, the signed distance correlation networks also obtain a higher score than their competitors constructed using Pearson correlation. The only exception to this trend is when comparing networks with the same average edge weight in the yeast dataset. However, overall, the presented results suggest that the networks constructed using signed distance correlation capture more biological information.

As shown in Table 5, our weighted and thresholded networks capture more biological information than unweighted networks. We extract the same conclusion from the analysis of the three datasets. This result is in line with what we expect since the use of edge weights representing the strength of the correlation between the expression of the pairs of genes results in an increase in the amount of information contained in the network.

The threshold values that we select to generate the weighted and thresholded networks from the three datasets are very similar to those that we use to construct the unweighted networks. This fact suggests that independently of the construction methodology we follow, the barrier between signal and noise is in the same range of correlation values for a given correlation matrix.

We use our method to construct a novel weighted and thresholded gene coexpression network for *R. leguminosarum*. This network promises to reveal rich biological information and it is therefore the starting point for further investigations of the biological mechanisms of this organism. In particular, we plan to identify groups of genes in *R. leguminosarum* which are highly connected in the network and associate them with specific biological processes. To do so, we will make use of community detection techniques and new experimental data. For the human liver dataset one could similarly validate communities by explore predicting disease-related biological information; see for example [20, 5, 14].

Finally, we have showcased our method on gene expression datasets from different organisms obtained using different techniques: microarrays (*R. leguminosarum*), RNA-Seq (yeast), and single-cell RNA-Seq (human). However, the methods that we have developed are general, and can also be used to construct networks in a vast range of domains, such as, for example, economics [26], neuroscience [2], climatology [6], and indeed any discipline where networks are constructed from correlation data.

## Acknowledgements

We thank Lyuba V. Bozhilova for her previous work on which this publication is built. We thank Alison K East and Florian Klimm for help and advice on how to analyse the datasets used in this manuscript. We thank Katharine Turner (ANU), Lyuba V. Bozhilova, Florian Klimm, and Malte D Luecken for fruitful discussions. In addition, we acknowledge support from COST Action CA15109, Keble College Oxford and Keble Association. The authors would like to acknowledge the use of the University of Oxford Advanced Research Computing (ARC) facility in carrying out this work (http://dx.doi.org/10.5281/zenodo.22558).

## Funding

This work is supported by the Engineering and Physical Sciences Research Council (EPSRC) [EP/R512333/1 to JPD, MBD, PSP, CMD and GR], the Biotechnology and Biological Sciences Research Council (BBSRC) [BB/T001801/1 to PSP and GR], the COSTNET COST Action [CA15109 to GR], and e-Therapeutics plc. MBD acknowledges support from the Oxford-Emirates Data Science Lab.

## Conflict of Interest

None

## Notes

https://github.com/javier-pardodiaz/sdcorGCN

### Competing Interest Statement

The authors have declared no competing interest.

